# Somite surface tension buffers imprecise segment lengths to ensure left-right symmetry

**DOI:** 10.1101/2020.08.14.251645

**Authors:** Sundar R. Naganathan, Marko Popović, Andrew C. Oates

## Abstract

The body axis of vertebrate embryos is periodically segmented into bilaterally symmetric pairs of somites. The anteroposterior (AP) length of somites, their position and left-right symmetry are thought to be molecularly determined prior to somite morphogenesis. Here we discover that in zebrafish embryos, initial somite AP lengths and positions are imprecise and consequently many somite pairs form left-right asymmetrically. Strikingly, these imprecisions are not left unchecked and we find that AP lengths adjust within an hour after somite formation, thereby increasing morphological symmetry. We find that AP length adjustments result entirely from changes in somite shape without change in somite volume, with changes in AP length being compensated by corresponding changes in mediolateral length. The AP adjustment mechanism is facilitated by somite surface tension, which we show by comparing *in vivo* experiments and *in vitro* single-somite explant cultures with a mechanical model. Length adjustment is inhibited by perturbation of Integrin and Fibronectin, consistent with their involvement in surface tension. In contrast, the adjustment mechanism is unaffected by perturbations to the segmentation clock, thus revealing a distinct process that determines morphological segment lengths. We propose that tissue surface tension provides a general mechanism to adjust shapes and ensure precision and symmetry of tissues in developing embryos.

---

Vertebrates are characterized by a left-right (LR) symmetric musculoskeletal system that emerges from bilateral somites during embryonic development. LR symmetry is vital for adult mechanical movements and a loss of symmetry is often associated with debilitating skeletal disorders such as scoliosis.^1,2^ Symmetry is often assumed to be a default state in somite formation,^3,4^ however, it remains unknown how robust somite shapes and sizes at the same position along the body axis emerge on the left and right sides of the embryo.

Somites are 3D multicellular units, typically with an outer epithelial layer surrounded by a fibronectin-rich extracellular matrix, that form by segmentation of the presomitic mesoderm (PSM).^5,6^ The AP length of somites and their LR symmetry is thought to be determined in the unsegmented PSM by genetic oscillations of a segmentation clock and downstream molecular prepatterns.^5-11^ While mechanical processes have also been associated with somite morphogenesis,^12–16^ their role in determining AP length and LR symmetry, if any, is not understood. In general, a quantitative study of bilateral symmetry in somites is lacking owing to the technical difficulty in following 3D somite morphogenesis simultaneously on the left and right sides of embryos.

To shed light on this problem, we performed multiview light-sheet microscopy of zebrafish embryos (Supplementary Movie 1) and developed a computational framework to perform map projection (Fig. 1A, Supplementary Movie 2), followed by automated segmentation of somite boundaries (Sup Fig. 1, 2, 3). This approach allowed us to follow LR somite morphogenesis in real time. We first quantified the AP length, *L_AP_*, of somites one to six and observed that the initial lengths, immediately after somite formation, was variable (Fig. 1B, Sup Fig. 4A; Coefficient of Variation, CV, 0.13; 95% CI [0.11,0.16]). To check whether the molecular prepatterns that are thought to set *L_AP_* can explain this variability, we measured interstripe distance in *mespb* gene expression stripes, which represent the first molecular indication of segment length in the anterior PSM.^17^ We observed the variability in *L_AP_* to be similar in magnitude to that of *mespb* segmental lengths (Fig. 1, D and E), suggesting that imprecision in *L_AP_* could be the consequence of a variable prepattern. Strikingly, within an hour after somite formation, *L_AP_* adjusted and the variability decreased (Fig. 1B; CV, 0.08 [0.07,0.09] at 1 hr; Sup Fig. 4). By comparing the initial and two-hr lengths, we identified 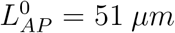 (Fig. 1C), which we defined as the target AP length, towards which somites tended to adjust. In other words, somites with 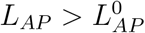 tended to become smaller and vice versa.

**Figure 1:**
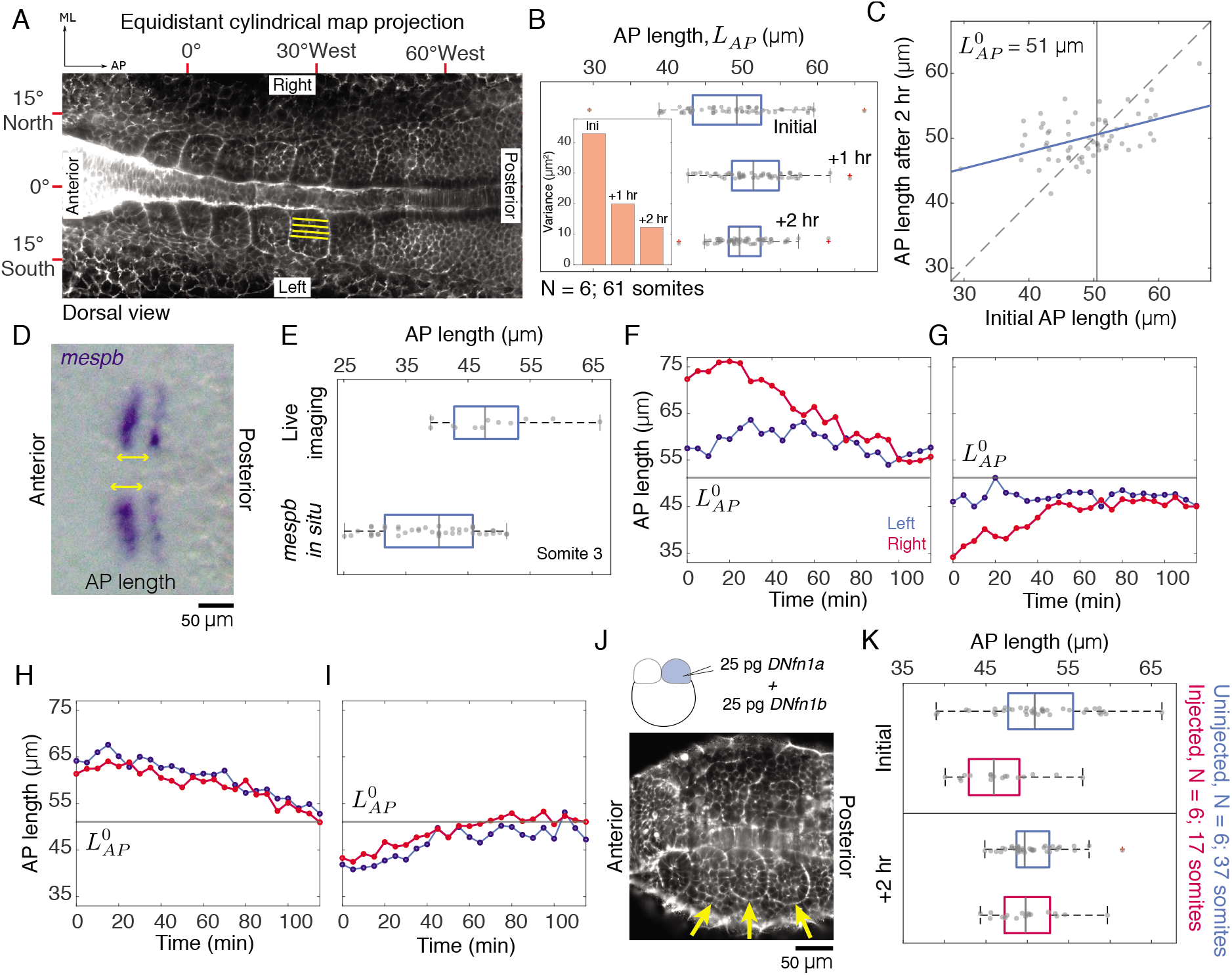
Initial somite lengths are variable and get adjusted independently on the leftright sides. (A) Map projected image of a 6-somite stage zebrafish embryo. AP, anteroposterior; ML, mediolateral; yellow lines, AP length (*L_AP_*) (B) Variability of *L_AP_* of first six somites decreases over time. Inset, variance of *L_AP_*. (C) Comparison of initial and two-hour *L_AP_*. Blue, linear regression; dashed line, slope=1; gray line, target AP length, 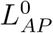 (D) Dorsal view of a flat-mounted embryo stained for *mespb* (blue). (E) Comparison of third somite *L_AP_* measured from live imaging and *mespb in situ* (arrows in (D)). (F-I) Representative plots of LR somite pairs, where only somites with initial *L_AP_* away from 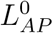 (gray line) adjust their lengths. (J) Schematic, DN fibronectin injection at 2-cell stage. Representative embryo with somite formation on one side (arrows) and no visible somites on the contralateral side. (K) Comparison of *L_AP_* of somites three-six in the somite-forming side of DN fibronectin injected embryos (red) and uninjected embryos (blue).

To investigate the mechanism of *L_AP_* adjustment, we first asked whether *L_AP_* on one side is influenced by lengths on the contralateral side. Comparing changes in length between corresponding LR somites, we observed that only somites with an initial length away from 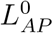 adjusted their lengths, regardless of the behavior of the segment on the contralateral side (Fig.1, F and G). In contrast, somites that formed with initial *L_AP_* close to 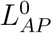 changed negligibly. Importantly, length changes occurred on the two sides only when initial *L_AP_* on both sides were away from 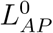 (Fig. 1, H and I). Combined, these results suggest that *L_AP_* changes on one side are not instructed by information from the contralateral side, but rather are determined by whether or not a particular somite has an initial length close to the target length.

We next asked whether the presence of any correctly formed somites on the contralateral side is required for length adjustments. To this end, we injected dominant negative (DN) *fibronectin 1a* mRNA together with DN *fibronectin 1b* in one of the cells at the 2-cell stage (Fig. 1J), which has been previously shown to perturb somite formation on one side.^18^ Injections resulted in 10% of the embryos bearing strongly disrupted somites on one side (Fig. 1J). When *L_AP_* of somites three to six was analyzed in the somite-forming side, we observed that lengths adjusted towards the same 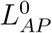 similar to uninjected embryos (Fig. 1K). While a possible cross talk mediated by fibronectin between the LR sides has been suggested,^19^ our results indicate that *L_AP_* adjustment on one side does not require somite morphogenesis in the contralateral side.

We next asked whether *L_AP_* adjustment is accompanied by a change in cell number or somite volume in the first hour after somite formation, when the majority of adjustment occurs. We observed that cell numbers increased in all somites (Sup Fig. 5B) irrespective of whether the initial *L_AP_* decreased or increased towards 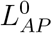. Accordingly, there was no significant correlation (r, −0.18 [-0.47,0.1]) between change in cell number and change in *L_AP_* (Fig. 2A). We next observed that somites exhibited a negligible change in volume (Fig. 2, B and C). We therefore conclude that neither a change in cell number nor change in somite volume mediates *L_AP_* adjustment.

**Figure 2:**
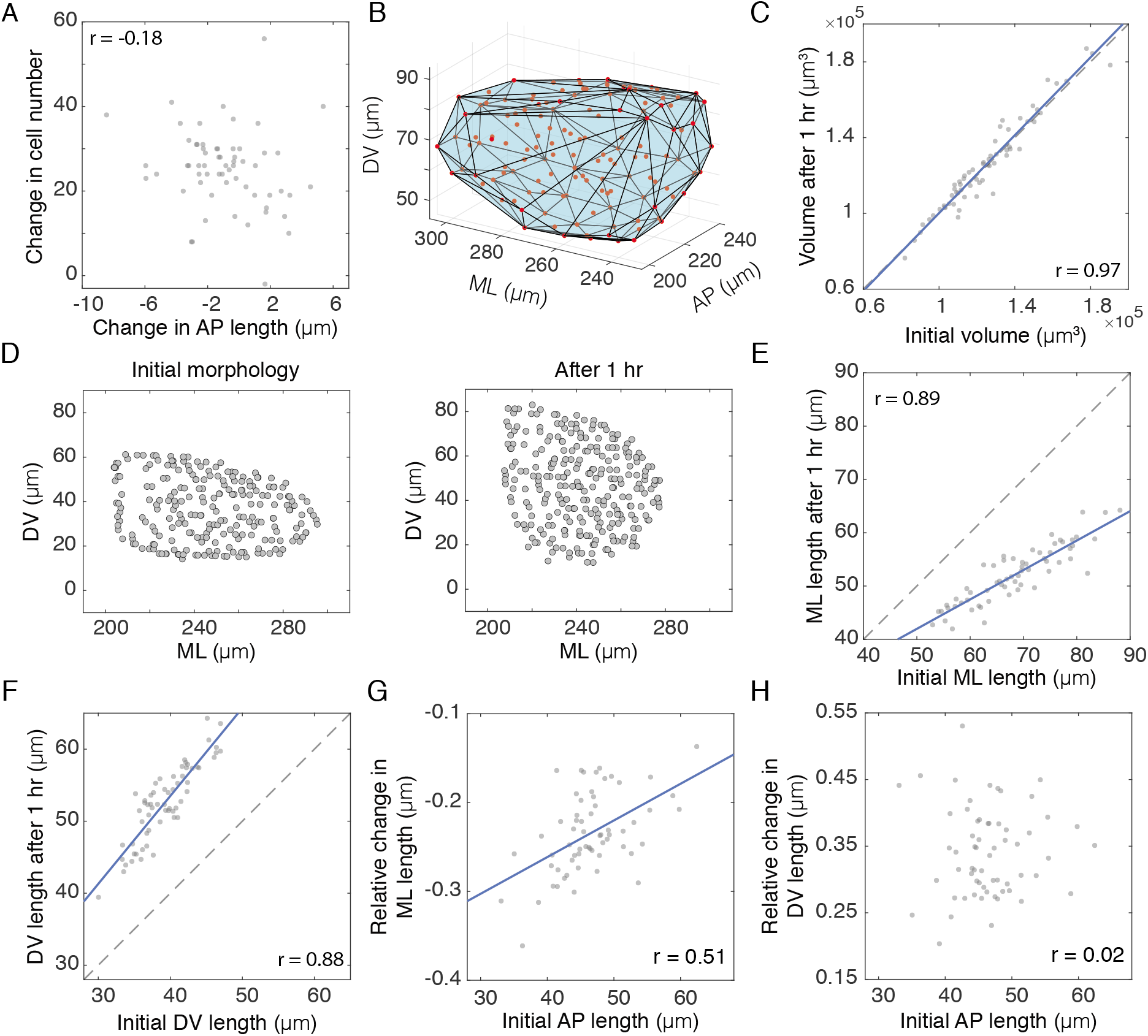
AP length adjustment results from somite shape deformations. (A) Comparison of change in AP length and change in cell number. (B) Convex hull of segmented nuclei (red) from a single somite. (C) Comparison of initial and one-hour somite volumes (r, 0.97 [0.95,0.98]). Blue, linear regression in (C, E-G) and dashed line, slope=1 in (C, E, F). (D) Nuclei (circles) distribution of a somite undergoing convergence-extension. (E-F) Comparison of initial and one-hour ML and DV lengths indicate decrease (r, 0.89 [0.82,0.94) and increase (r, 0.88 [0.82,0.93]) in lengths respectively. (G-H) Initial AP lengths are positively correlated with relative changes in ML length (*L_ML_*(1*hr*) — *L_ML_*(0*hr*))/*L_ML_*(0*hr*) (G), and not correlated with relative changes in DV length (*L_DV_*(1*hr*) — *L_DV_*(0*hr*))/*L_DV_*(0*hr*) (H).

This volume conservation constraint suggests that changes in *L_AP_* must be reflected in corresponding changes in the other two dimensions of the somite. We therefore quantified 3D shape changes (Materials and Methods) and observed that mediolateral (ML) somite lengths decreased over time, while dorsoventral (DV) lengths increased, reflecting convergence-extension (CE) in somites (Fig. 2, D-F, Sup Fig. 5). The initial *L_AP_* was positively correlated (r, 0.51 [0.27,0.68]) with relative changes in ML length (Fig. 2G). Thus, somites with an initial *L_AP_* smaller than 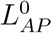 tended to have an increased reduction of ML length and vice versa. In contrast, initial *L_AP_* was not correlated (r, 0.02 [-0.25,0.3]) with relative changes in DV length (Fig. 2H) suggesting that DV length dynamics does not contribute to *L_AP_* adjustment. We conclude that *L_AP_* is adjusted by corresponding changes in ML length of somites implying that the ML dimension buffers imprecisions in AP length of somites. Importantly, these observations suggest that AP length robustness is associated with mechanical forces that drive somite shape changes.

To understand the role of mechanical forces in *L_AP_* adjustment, we next sought to develop a coarse-grained mechanical model of a newly formed somite, which we represent as a cuboid of constant volume (Fig. 3D(a)). Mechanical stresses acting on the somite consist of somite surface tension stemming from both extracellular matrix and somite epithelial cells, contact stresses with surrounding tissues and internal active stresses driving CE flows. How these stresses lead to somite shape changes is determined by the somite material properties, which we investigated by developing a single-somite explant culture (Fig. 3A, Methods). We carefully isolated somite three or somite four from embryos and followed their change in shape over several hours. Interestingly, all explanted somites (N = 5) became spherical over time (Fig. 3B, Sup Fig. 6, Supplementary Movie 3) in the absence of neighbouring tissues. This final spherical shape suggests that organisation of active CE flows in a somite is lost when explanted. Although the anterior PSM at later stages has been reported to behave as a yield stress material,^20^ in our experiments, the explanted somite behaves as a viscous fluid with surface tension (Fig. 3D(b)).

**Figure 3:**
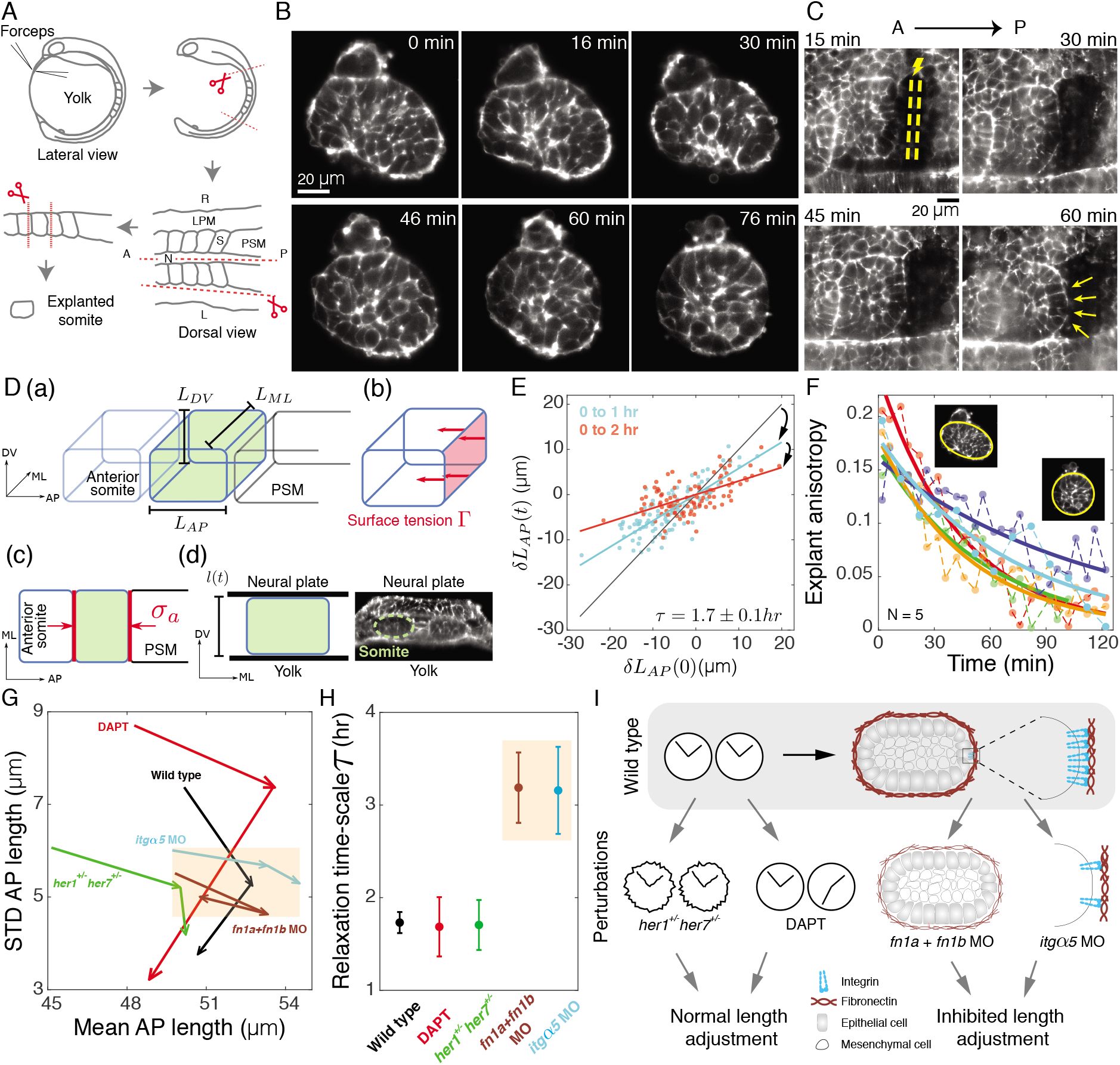
Somite surface tension facilitates length adjustment. (A) Schematic of somite explant preparation. LPM, lateral plate mesoderm; S, somite; N, notochord. (B) Explanted somite rounds up over time. (C) Ablation (yellow) of PSM adjacent to recently formed somite boundary. Yellow arrow, bulging of boundary. (D) (a) Schematic of somite dimensions. (b) Normal stress on somite surface due to surface tension. (c) Contacts (red) with PSM and anterior somite result in normal stress *σ_a_*. (d) Left, constraint *l(t)* imposed on *L_DV_* by neural plate and yolk. Right, snapshot of somite 3 from a 3-somite stage embryo. (E) AP length adjustment is proportional to the initial variation from 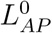, consistent with model prediction. Linear fit of the data (blue, 0 to 1 hr; red, 0 to 2 hr) yields the relaxation time-scale *τ*. Gray line with slope=1 is a reference for no change in *L_AP_*. (F) Shape anisotropy of 5 explanted somites over time. Insets, representative initial and final shape. (G) Phase diagram of change in standard deviation of *L_AP_* over 2 hrs. For each condition, the two arrows represent 0 to 1 hr and 1 to 2 hr transition respectively. Orange box highlights reduced AP adjustment (G) and longer relaxation time-scales (H) due to perturbation of molecules involved in surface tension. (H) Estimated relaxation time-scale *τ* for the different perturbations. Error bar, fit uncertainty. (I) Schematic of effect of different perturbations on somite length adjustment.

To investigate contact stresses in the AP direction, we performed laser ablation of the PSM posterior to the most recently formed somite boundary. We observed that over time a bulge of the somite boundary next to the ablated site appeared, indicating that a compressive normal stress exists between the PSM and the somite (N = 6, Fig. 3C, Sup Fig. 7). This is consistent with previous experiments in chick^12^ and later-stage zebrafish embryos.^21^ We include this stress in the model as normal stress *σ_a_*(*t*) acting on both the anterior and posterior surfaces of the somite (Fig. 3D(c)). Along DV dimension, somites are sandwiched between neural plate and yolk at early stages, imposing a constraint *l*(*t*) on the DV extension of the somite (Fig. 3D(d)). Finally, we account for CE flows in the model through an internal active shear stress.^22,23^ For simplicity, we do not specifically account for contact stresses along ML dimension (Supplementary Information) and we neglect frictional forces between somites and surrounding tissues. By considering a linear viscous fluid model of somite tissue, together with these boundary conditions, we obtain a dynamical equation for *L_AP_* (Supplementary Information Eq. 18). We find that 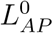 is determined by a combination of surface tension, external stresses and CE flows (Supplementary Information Eq. 19). Furthermore, assuming a constant 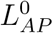, we find that variations of somite AP length from the target value 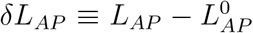 relax in time, following approximately

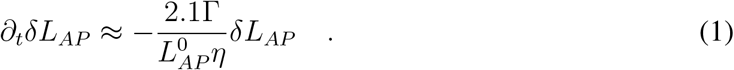

Here, *η* is the somite viscosity and 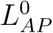 is determined by *σ_a_*(*t*), *l*(*t*), internal active stresses and surface tension Γ. Therefore, variations of *L_AP_* are reduced in time over the time-scale 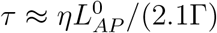. Eq. 1 predicts *L_AP_*(*t*) to be proportional with *L_AP_*(*0*), with proportionality coefficient exp(—t|τ). Consistent with this, we find that changes in *L_AP_ in vivo* are proportional to their initial values (Fig. 3E), which allows us to extract the relaxation time-scale *τ* = 1.7 ± 0.1 hr using a linear fit to the data (see Supplementary Information). In order to estimate the relevance of surface tension, we then quantified relaxation of explanted somites towards a spherical shape, which is driven only by surface tension (Fig. 3F, Sup Fig. 6, Supplementary Information). We find the relaxation time-scale *τ*_e_ to be 1.1 ± 0.4 hr. Since *τ*_e_ is comparable to *τ* (Supplementary Information), we conclude that for a given 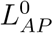, stresses generated by surface tension can account for a major part of the observed adjustment.

If tissue surface tension is indeed critical for length adjustments, then we expect adjustments to change when surface tension is perturbed. Tissue surface tension is known to be determined by a combination of cell-ECM interaction, cell-cell adhesion and actomyosin activity.^24-26^ We therefore targeted the Fibronectin-rich ECM by injecting morpholinos (MO) against both *fibronectin1a* and *1b* in 1-cell stage embryos. MO injection above 1.5 ng caused anterior somites to disintegrate by the 10^th^ somite stage (Sup Fig. 9B) consistent with previously published results.^27^ However, at lower injected amounts (about 1 ng), anterior somites remained intact and when we quantified AP lengths of somites 2 to 6, we observed that the length adjustment over 2 hrs was strongly reduced (Fig. 3G, Sup Fig. 9, A and B). We obtained a similar result by targeting cell-ECM interaction through MOs against *integrinα5* (Fig. 3G, Sup Fig. 9, A and B). We quantified the relaxation time-scale *τ* under both these conditions and observed a significant increase compared to wild type embryos (Fig. 3H). These results show that perturbing molecules implicated in tissue surface tension reduces somite length adjustment (Fig. 3I).

We next wondered if length adjustment still occurs after mild perturbations to the segmentation clock in which somite boundaries are not defective. We first targeted the core clock circuit through heterozygous *her1;her7* mutants and observed that mean initial somite length was shorter, yet somite length variability still reduced within 2 hrs similar to wildtype (Fig. 3G, Sup Fig. 9A). We then targeted the Delta-Notch signaling pathway by treating embryos with DAPT, which is known to desynchronize the clock and cause defects restricted to boundaries posterior to segment 6.^28^ We observed that the initial variability in lengths of somites 2 to 6 was higher, potentially reflecting elevated noise in the prepatterning process, but still these lengths adjusted normally (Fig. 3G, Sup Fig. 9A). Importantly, in both cases the relaxation time-scale *τ* was similar to wild type embryos (Fig. 3H). Taken together, these results suggest that the adjustment mechanism is distinct from the clock (Fig. 3I).

So far, we have described the mechanism of *L_AP_* adjustment from a unilateral perspective. What is the consequence of these unilateral length adjustments for the bilateral symmetry of somites? We reasoned that if both the posterior boundary of the head mesoderm^29^ and cell flow into the PSM are LR symmetric,^30^ length adjustments would simultaneously ensure LR symmetrical somite lengths and segment boundary positions along the body axis. We observed that initial AP length differences were variable (CV, 0.11 [0.09,0.12]; Fig. 4, A and B), as were the boundary position differences (CV, 0.13 [0.1,0.15]; Fig. 4, B and C), with a lack of bias between left and right sides, indicating that many bilateral pairs form asymmetrically. However, as somite lengths adjusted, both length differences (CV, 0.07 [0.06,0.08]) and boundary position differences decreased (CV, 0.09 [0.08,0.1]), leading to a more symmetric segmented morphology. Although the initial length difference was only weakly correlated with anterior boundary position difference (r, −0.24 [-0.45,-0.03]) (Sup Fig. 10B), it correlated significantly with the posterior boundary position difference (r, 0.67 [0.51,0.8]) (Fig. 4D). Combined, these findings indicate that as new somite pairs form, their anterior boundaries are symmetric, either because they were initially symmetric or because they had substantially adjusted, and so any asymmetries in length are predominantly a result of asymmetric positioning of the most recently formed boundary. A change in position of the posterior boundary ensues from length adjustment, simultaneously leading to increased LR symmetry in both AP lengths and boundary positions (Fig. 4, E and F). Overall, our results show that unilateral length adjustments facilitated by somite surface tension ensures increased precision and bilateral morphological symmetry of somites.

**Figure 4:**
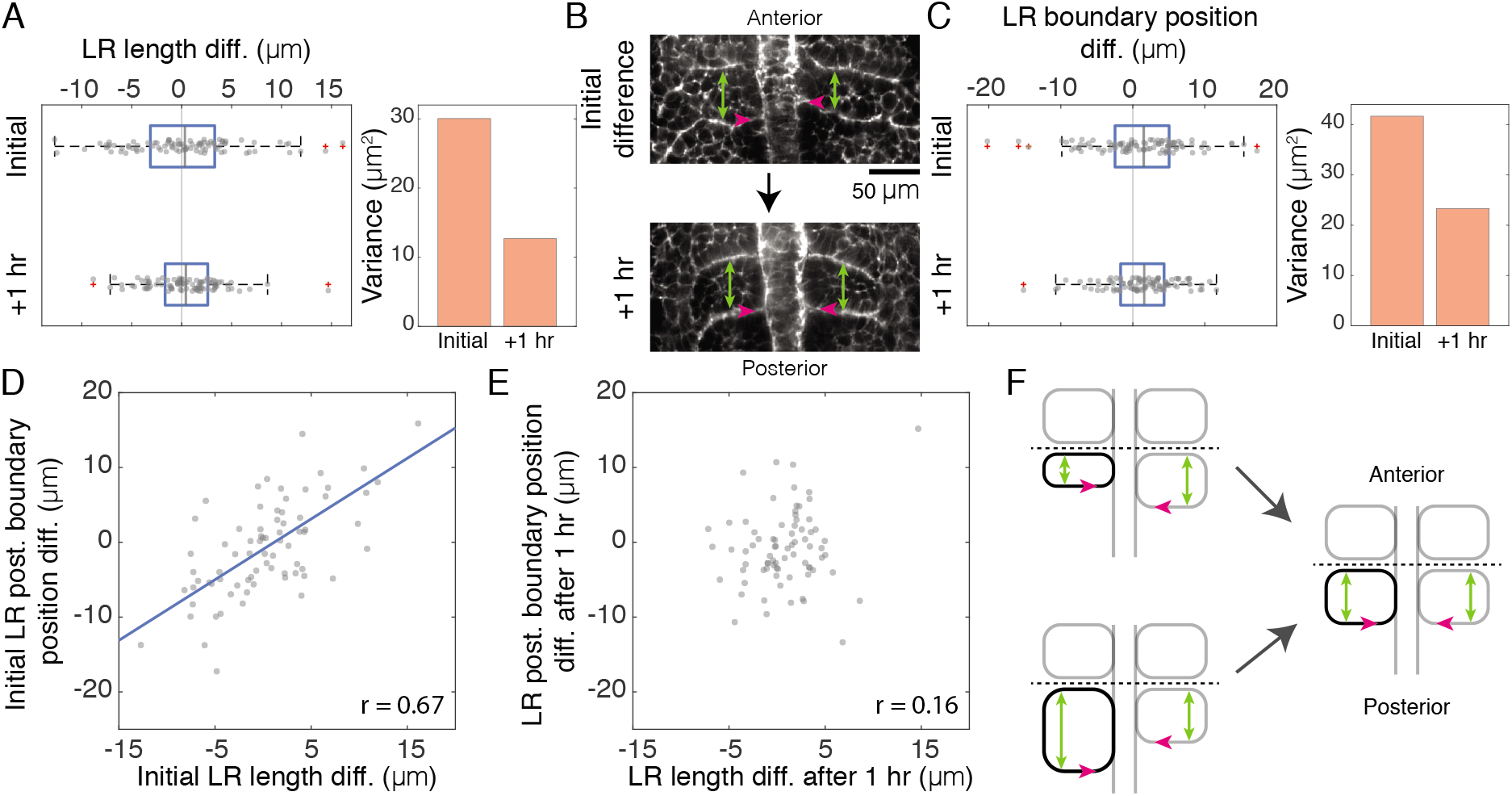
Somite lengths and boundary positions increase symmetry over time. (A) Variability in AP length difference between left-right somite pairs decreases over time. Left, box plot; Right, variance. (B) Representative images of left-right somite pairs with asymmetric initial lengths (green) and somite boundary positions (pink) that adjust over time. (C) Variability in left-right boundary position difference decreases over time. Left, box plot; Right, variance. (D-E) AP length difference between left-right somite pairs is positively correlated initially with posterior boundary position difference (D), while no significant correlation is observed (r, 0.16 [-0.19,0.48]) after 1 hour (E). Blue, linear regression. (F) Schematic of initial asymmetries in somite length (green) and position (pink) adjusting concurrently. Dashed line represents aligned anterior boundaries.

The AP length of somites has been historically understood from the perspective of the segmentation clock and downstream molecular processes in the PSM, and bilateral somite formation has largely been considered as symmetric.^3^ Asymmetry was thought to arise only when retinoic acid signalling was lost, exposing molecular prepatterns in the PSM to a gene expression program that determines left-sided organ positioning.^9-11,31^ However, our findings that initial lengths are imprecise, but are adjusted by 3D somite deformations, show that this perspective is insufficient to describe the length and symmetry of somites. In addition to the prepattern, we argue that somite surface tension, external stresses from neighboring tissues and CE flows within somites must also be included. Similar to the hypothesis proposed in,^32^ our results suggest that the role of the prepattern is to provide a coarse allocation of material for each somite, which is then fine-tuned by tissue mechanics. The LR differences in somite formation observed here, in otherwise normally-developing wild type genetic backgrounds and in constant environmental conditions, reawakens the idea that links subtle developmental failures in left-right symmetry to idiopathic scoliosis in humans.^33^

Bilateral symmetry is a feature of many organ systems (e.g. eyes, ears and kidneys). Similar to somites, symmetry in bilateral ears also emerges over time, but in this case through differential growth between the LR ears triggered by lumenal pressure.^34^ A role for mechanics in symmetrization of body plans extends beyond the bilateria, as demonstrated by the action of muscle contraction in recovery of radial symmtery in Cndaria.^35^ Our work showing how tissue surface tension ensures precision of somite morphology joins recent studies of mechanical processes reported to buffer heterogeneous cell growth in sepals in plants^36^ and to enable straight cephalic furrow formation in *Drosophila* embryonic epithelia,^37^ revealing mechanics as a general principle in ensuring developmental precision.

Since the function of a tissue is intimately related to its form and shape, this newly identified role of mechanics in controlling the precision of tissue shape has implications that go beyond developmental patterning to other fields where a precise final tissue shape is critical. In tissue engineering and regenerative medicine, where recent applications using organoids and other *ex vivo* analogs of developmental tissue strive for reproducible shapes,^38^ our findings suggest that understanding how mechanics contributes to precision in these settings could help to over-come current limitations. Our findings also raise the possibility that during the evolution of new developmental patterns via mutations to the underlying genetic regulatory networks,^39,40^ resulting fluctuations in morphology may be stabilized by tissue mechanics, potentially facilitating a greater search of sequence space while maintaining a precise body architecture.

## Materials and Methods

### Zebrafish care

Wildtype (AB), transgenic and mutant fishes were maintained according to standard procedures and all embryos were obtained by natural spawning. Utr::mCherry transgenic line (e119Tg), originally established in the Heisenberg lab was obtained from the Mosimann lab. H2B::GFP transgenic line (kca6Tg) originally established in the Campos-Ortega lab was obtained from the MPI-CBG fish facility. Heterozygous animals from these transgenic lines were used for crossing. Homozygous *her1;her7* mutant fishes were crossed with the heterozygous Utr::mCherry transgenic line to generate heterzygous mutants. Immediately after fertilization, embryos were shifted to a 33°C incubator and grown to developmental stages of interest before time-lapse imaging. All experiments were carried out using embryos derived from freely mating adults, and thus are covered under the general animal experiment license of the EPFL granted by the Service de la Consommation et des Affaires Vétérinaires of the canton of Vaud - Switzerland (authorization number VD-H23).

### Multiview imaging

Polytetrafluoroethylene (PTFE) tubes (Adtech, Part No: STW15, Lot No: D22869; inner diameter, 1.58 mm), were first cleaned as described^41^ and pre-cut to 2.5 cm. The cut tubes were then straightened by heating in an eppendorf with water at 70°C for 5 minutes. Zebrafish embryos in their chorion between 50% and 75% epiboly were then transferred to 0.25% low-melting agarose solution prepared with E3 fish medium. The agarose solution in addition contained 0.5 μm green fluorescent beads (Thermo Scientific, Lot No: 172285), which were used for image registration. For 10 ml of agarose solution, 4 μl of bead solution was added and mixed thoroughly. Before solidification of agarose, two embryos were loaded into each tube. 15 PTFE tubes were loaded with embryos and kept upright in an eppendorf filled with E3 medium. The embryos were allowed to grow until the 1-somite stage in the PTFE tubes at 33°C. Using bright-field illumination in a Zeiss Z1 light-sheet system, an embryo with its notochord approximately along the circumference of the tube was then chosen for time-lapse imaging (Sup Fig. 2A).

Transgenic embryos with fluorescent markers for visualisation of actin filaments and nuclei (Utr::mCherry;H2B::GFP transgenic line) were imaged from 6 angles (30° apart, Sup Fig. 2B) for 6 hours from the one-somite stage at 28°C. A 488 nm laser (25% power, 100 ms exposure) and a 561 nm laser (50% power, 100 ms exposure) with light-sheet thickness of 4.5 μm, along with two bandpass emission filters (BP 505-545 and BP 575-615) were used for imaging the two fluorophores. Multiple z-slices (between 80 and 100; 2 μm apart) were acquired in each angle at a time interval of 5 min. Before the start of the time-lapse, the PTFE tube was translated along the y-axis and fluorescent beads were imaged from the 6 different angles. A 20x/1 NA detection objective, two 5x/0.16 NA illumination objectives and a PCO Edge 5.5 camera with a pixel size of 6.5 μm by 6.5 μm were used for imaging.

### Processing multiview movies

Using the FIJI^42^ Multiview reconstruction plugin^43^ and fluorescent beads as registration markers, initial registration of the six angles was performed using rotation invariant matching. This transformation was then applied to the embryo images. Nuclei were then used as markers to perform two further rounds of registration to correct fine drifts between time points. For this, the translation invariant transformation was first applied following which a precise matching was performed using the iterative closest point algorithm. The transformation obtained from the nuclei channel was then applied to the Utr channel. The point spread function for the imaging system was then determined from the bead images. Following a successful registration, fusion (Sup Fig. 2C, representative fused image) of the different angles with deconvolution was performed after downsampling by a factor of two. Default parameters from the multiview reconstruction plugin were used for each of the aforementioned steps.

Custom MATLAB algorithms were developed for further processing of the fused images as described below.

#### Nuclei segmentation

The downsampled fused image had a pixel size of 0.46 μm. To detect nuclei, the laplacian of gaussian filter (with a filter size of 15 pixels and standard deviation of 5 pixels) was applied on every 5^th^ z-slice of the fused image and local maxima (defined as spots) that represented potential nuclei positions were determined. The mean and standard deviation (SD) of spot fluorescence intensities were determined and only those spots with a fluorescence intensity higher than one SD were considered for further processing. The local signal to noise ratio (SNR) was then determined in a box of size 9 by 9 pixels surrounding spots of interest in contrast to a local region (10 by 10 pixels) surrounding each of the boxes. We observed that a SNR greater than 2 signified nuclei positions. The nuclei thus identified in every 5^th^ slice was used to generate a 3D point cloud to which a sphere fit was performed (Sup Fig. 2D) using linear least squares with the ellipsoid fit function (developed by Yury Petrov, MATLAB Central File Exchange). The radius of the sphere did not change significantly during the analysis period (Sup Fig. 2E). The centre and radius of the sphere were then used for performing map projection of the fused images.

#### Equirectangular cylindrical map projection

For each fused image, a multi-layered cylindrical projection was performed by generating 80 to 100 concentric circles with a step size of 2 μm around the estimated radius. This allowed for unwrapping different layers of the 3D fused embryo onto 2D surfaces. For each layer of the fused embryo, an empty map was first generated that extended from −35° to 35° in the y-direction and 0° to 170° in the x-direction. Using equirectangular cylindrical projection formulas, the latitudes and longitudes that correspond to each position in the empty projected map were then determined. The cartesian (x,y,z) positions that correspond to each of these (latitude, longitude) points in the projected map was obtained using standard spherical to cartesian coordinate system conversion formulas. Following these transformations, a direct mapping of the pixel values corresponding to a (x,y,z) position in the fused image onto the projected map was performed.

A scale factor of 15 was used for the projection, which resulted in a pixel size of about 0.35 μm along the equator for a map generated with the estimated radius. Note that the pixel size changes away from the equator in a single map as well as across maps generated with different radii. Because embryos were oriented with their notochord along the circumference of the tube, somites formed within 15° of the equator in the projected maps (Fig. 1A), where negligible distortion occurs. In a map of a particular radius, the change in pixel size from the equator to 15° is less than 0.05 μm. Similarly the change in equatorial pixel size for different layers of the projected maps, where somites could be visualized, was less than 0.1 μm. Thus, when compared to the target AP length (51 μm), the change in pixel size both across maps and within a map was negligible. Therefore, for quantifications of somite characteristics, an average pixel size was used, which was determined across the different map projection layers.

#### Somite boundary and notochord segmentation

In the map projected images, the notochord was horizontal and somite boundaries more or less orthogonal to the notochord. Given this distinction, the images were first subdivided into two parts, one that contained all lines that were approximately horizontal (90 ± 30°) and one that contained all other lines. To perform this, the Frangi vesselness filter^44^ was employed on 2D images, which computes the likeliness of an image region to vessels or tube-like structures by computing the eigenvectors of the Hessian of the image. This filter was downloaded from MATLAB Central File Exchange (Hessian based Frangi Vesselness filter developed by Dirk-Jan Kroon). The following settings were used in the filter: ‘Frangiscalerange’, [1 3], ‘FrangiScaleRatio’, 2, ‘FrangiBetaOne’, 0.5 and ‘FrangiBetaTwo’, 12. The direction of the eigen vectors was then used (Sup Fig. 3, C and G) to subdivide the image into two parts.

The sub-image that contained horizontal lines i.e the image with the notochord was first considered. Along each column of the image, the *findpeaks* function from MATLAB was employed (with a minimum peak height that was at least one SD above the mean fluorescence intensity along the column) to detect local increase in fluorescence intensities. Note that along the notochord as well as along somite boundaries, an increase in fluorescence intensity is observed. Upon applying this function, all pixels with a threshold increase in fluorescence intensity were obtained, following which those identified pixels that were less than 15 pixels apart were joined by a straight line. This resulted in formation of continuous horizontal lines in the image and the notochord was then manually identified by choosing lines of interest and by deleting all other horizontal lines (Sup Fig. 3, D-F). The same methodology was employed along each row in the sub-image that contained non-horizontal lines i.e the image with somite boundaries (Sup Fig. 3, H-J). For one of the time frames, this segmentation was performed in all map projected layers and the segmented lines were projected on top of each other. We observed that the segmented lines of a single boundary spread over a width of 2 μm across projected layers. Given such a small deviation in boundary position across map projected layers, segmentation was performed only on single map projected layers for all subsequent analysis.

#### Somite length and boundary position quantification

The analysis was started only upon formation of the first two boundaries i.e upon formation of the first morphological somite in the embryo. A boundary was said to be formed when an increase in utrophin fluorescence intensity was observed along the entire mediolateral extent. Upon formation of a boundary, the (x,y) coordinates of the segmented notochord closest to the somite boundary under consideration was noted. Similarly, for the immediate anterior boundary, the (x,y) coordinates of the notochord closest to the boundary was noted. The local angle of the notochord was then determined between these two notochord positions. Prior to somite length and boundary position determination, the first 15 pixels (about 6 μm) of the somite boundary closest to the notochord were ignored as they were found to be noisy in the images. A line with the determined local notochord angle was then drawn along each of the subsequent 60 pixels (about 25 μm) from one boundary to the next and the average somite length was determined (Sup Fig. 3B). Somite lengths from the left were subtracted from the right sides thus providing a length difference. To determine boundary position, a line perpendicular to the local notochord angle was drawn from each of the 60 pixels of interest along a boundary and the point of intersection of these lines with the notochord were noted (Sup Fig. 10A). The median position across these points of intersection was taken as the boundary position. A difference in boundary position between the two sides informed on whether a particular boundary was more anterior or posterior when compared to its contralateral pair.

### Single-view light-sheet imaging

Viventis microscope was used for performing single-view light-sheet imaging, which allowed for imaging multiple (four to six) embryos simultaneously. The embryos were dechorionated and carefully placed dorsally in a 3D-printed imaging chamber filled with 600 μl E3 fish medium. The imaging chamber had troughs with a width of 650 μm. Fluorescence excitation was achieved with a dual illumination scanned Gaussian beam light-sheet of appox. 3.3 μm thickness (full width at half maximum) using 488 and 561 nm lasers. The signal was collected with a Nikon CFI75 Apo LWD 25x/1.1 NA objective and through 525/50-25 and 561/25 nm bandpass filters respectively onto an Andor Zyla 4.2 Plus sCMOS camera. The microscope is equipped with an incubation chamber to maintain the embryos at 28^°^C. 70 z-slices with a spacing of 2 μm and a frame interval of 5 min were acquired for 4 hrs. The pixel size under these settings was 0.35 μm. Embryos were imaged from the one to six somite stage using the Viventis system. Single-view data set for wild type embryos were used for cell number, volume quantification and for comparison of AP lengths with ML and DV lengths (Fig. 2, Sup Fig. 5). For the linear data fit of AP length relaxation (Fig. 3E) as well as for analysis of LR symmetries in somite length and boundary position (Fig. 4, Sup Fig. 10), multiview data set from Zeiss and single-view data sets were pooled.

### Cell number and volume quantification

To quantify cell numbers in somites from Viventis movies, the images were first converted to xml/hdf5 format using BigDataViewer FIJI plugin.^45^ Mastodon FIJI plugin (Version 1.0.0-beta-17, https://github.com/mastodon-sc/mastodon) was then used for nuclei segmentation. The utrophin channel was first selected and a region of interest that represented a somite was first specified. A radius of 5 μm and a quality factor of 25 was then used for nuclei segmentation in the histone channel. A manual correction was then performed to remove nuclei that were detected outside a somite and to add nuclei that were missed in the segmentation process. The data set was then transferred to MATLAB and a convex hull was applied on the detected nuclei (Fig. 2B) to determine the volume of a somite. We expect an error of 5 to 10 cells in the cell number quantification and this error is predominantly due to segmentation errors in the ventral most part of somites, which has lesser contrast due to increased scattering of laser light deeper in the embryo.

### Quantification of 3D somite dimensions from nuclei positions

Given the segmented 3D nuclei distribution in somites we adjusted the coordinate system to coincide with the main embryo axes. During data acquisition somites three to five were oriented such that they were positioned in the same focal plane. However, occasionally somites are slightly tilted in ML-DV and ML-AP planes. To account for this, we rotated left-right somite pairs first around the AP axis so that individual somite centres of mass lie on the same DV position, and then around the DV axis so that the centres lie on the same AP position.

We then used the somite nuclei positions to define the somite shape tensor Q as

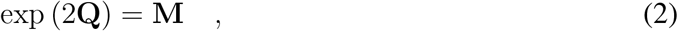

where M is moment of inertia tensor of the cell nuclei distribution. Q quantifies the strain required to deform an isotropic shape into the shape of the somite, similar to shape tensors defined in^22^ to describe 2D shape of *Drosophila* wing. We then defined AP, ML and DV dimensions of somites as

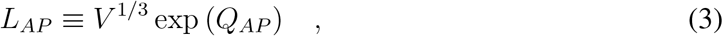

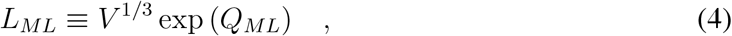

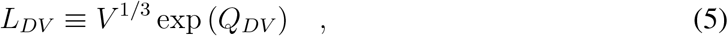

where *Q_AP_,Q_ML_,Q_DV_* are the corresponding diagonal elements of the somite shape tensor **Q**.

As expected, this definition yields lower values of *L_AP_* than the direct boundary segmentation since it is based on nuclei positions. However, we find that the two measures are well correlated (r = 0.8), as shown in Sup Fig. 5C. We use these definitions only when discussing the ML and DV dimensions in comparison to the AP and the boundary segmentation method is employed otherwise.

### Statistics

The coefficient of variation (CV) for AP lengths was simply determined by dividing standard deviation by the mean. For length differences and boundary position differences, the standard deviation was divided by the target AP length (51 μm) to estimate CV. For determining confidence intervals (CI) for CV, data set of interest was subjected to the built-in bootstrap function in MATLAB with 100,000 repeats with replacement. For determining CI for correlation coefficients, pairs of variables were subjected to bootstrap. In all box plots, left and right edges of rectangular box represent the 25^th^ and 75^th^ percentiles respectively, gray line represents the median, whiskers extend to 1.5x the interquartile range and plus symbol represents outliers.

### *In situ* hybridization

*mespb* riboprobe was generated from a plasmid.^46^ Utr::mCherrry;H2B::GFP double transgenic embryos were fixed at the 2-somite stage in freshly prepared 4% paraformaldehyde solution overnight. *In situ* was then performed in a 24-well plate as described.^47^ Following staining, embryos were flat-mounted in glycerol and documented with a stereomicroscope (Olympus SZ61) equipped with a digital camera (DP22, Olympus). The segment lengths between *mespb* stripes were determined manually close to the notochord in FIJI.

### Embryo injection

N-terminal 70 KDa fragments of dominant negative *fibronectin 1a (DNfn1a)* and *fibronectin 1b (DNfn1b)* plasmids were obtained from the Heisenberg lab. mMESSAGE mMACHINE Kit from Life Technologies was used for performing *in vitro* transcription of these plasmids using the sp6 primer. 25 pg of *DNfn1a* together with 25 pg of *DNfn1b* was injected in one of the cells at the 2-cell stage of embryos obtained from a cross between Utr::mCherry and H2B::GFP heterozygous lines. The injection solution with these constructs was prepared in RNAse-free water, which also contained 10% phenol red for visualization of injection with naked eye. Injection needles were prepared using a glass capillary (Model No. GC100F-15, 1.0 OD x 0.58 ID x 15 L cm) in a needle puller (WPI Sutter instrument, Model P-97) and were back filled with 3 μl of injection solution. A droplet of mineral oil (Sigma, M3516) was added to a microscope stage micrometer, 1 mm/0.01 mm scale (Cole-Parmer, Meiji Techno MA285, Item No. GZ-48404-8) and injection solution was injected into the mineral oil. The needles were manually snipped using forceps until injection resulted in a 100 μm sized droplet, which corresponded to an injection of 0.5 nL. The working concentration of the mRNAs were adjusted such that 0.5 nL injection corresponded to 25 pg of mRNA. Injection was performed using a Pneumatic Pico Pump PV 820 (World Precision Instruments), an eject pressure of 20 psi, and a little back-pressure so as to ensure no liquid in-or outflux of the needle between injections. About 100 embryos were injected and grown at 33°C until the 2-somite stage. Embryos with perturbed somites on one side of the body were then selected (about 10% of injected embryos exhibited this phenotype). Somites three to six in the selected embryos positive for both transgenes were used for imaging in the Viventis light-sheet system. Morpholinos for *itgα5* (5’-TAACCGATGTATCAAAATCCACTGC-3’), *fn1a* (5’-TTTTTTCACAGGTGCGATTGAACAC-3’) and *fn1b* (5’-GCTTCTGGCTTTGACTGTATTTCGG-3’) were obtained from Gene Tools, LLC, diluted in RNAse free water and injected at indicated concentrations in the cytoplasm of 1-cell stage embryos.

### Widefield imaging

Morpholino-injected embryos were first subjected to bright field imaging (Sup Fig. 9B) using the Zeiss Axio Observer 7 widefield microscope to ensure that overall embryo morphology as well as somite morphology are unaffected at injection amounts that were ultimately used for analysis of length adjustments. Embryos obtained from a cross between Utr::mCherry and H2B::GFP heterozygous lines were dechorionated at 1-somite stage and transferred to a multi-embryo imaging mold with E3 medium. Embryos were then imaged at 28°C using a Fluar 10x/0.5 NA M27 objective and Prime95B back-illuminated sCMOS camera with a frame interval of 10 min for 7 hrs.

### DAPT treatment

50 mM DAPT stock solution (Merck) was prepared in 100% DMSO (Sigma) and stored in a small volume at −20°C. A working solution of DAPT was prepared fresh before each experiment and the treatment was performed in 12-well plates. To prevent precipitation, the DAPT stock solution was serially diluted 4 times to reach the final working concentration of 25 μM. To a single well that contained 1.6 ml E3 medium with DAPT, 25 embryos in their chorions at shield stage were transferred. The embryos were allowed to grow until 1-somite stage in the 12-well plates at 33°C. 6 embryos positive for both utrophin and histone transgenes were then dechorionated and transferred to a Viventis imaging chamber containing 25 μM DAPT and time-lapse imaging was performed. DAPT was left in the chamber for the entire duration of imaging.

### Laser ablation

Custom-built UV laser ablation system in the EPFL microscope facility was used for ablating the presomitic mesoderm. The ablation set up is attached to a confocal spinning disk unit (Yokogawa CSU10B-F300-E). Embryos at the 2-somite stage obtained from a cross between Utr::mCherry and H2B::GFP heterozygous lines were dechorionated and mounted on their dorsal sides in a Viventis imaging chamber. PSM posterior to somite three or four was ablated in all experiments. Pulsed UV laser (50 mW, 355 nm, 0.1 ns at 1000 Hz, model PNV-M01510-130) at 80% power was used for the ablation. A 70 μm line oriented along the mediolateral axis immediately posterior to a most recently formed boundary was defined as the ablation region with the following settings: 1 point per micrometer and 5 pulses per point at maximum repetition (1000 Hz). Two ablations were performed, with the first ablation about 15 μm posterior to the most recently formed boundary and the second ablation about 10 μm posterior to the previous ablation. A N-Achromat 63x/0.9 CORR objective was used for ablation and bright field images before and after ablation were obtained using a CoolSNAP HQ air cooled CCD Camera with binning of 2. The imaging chamber was then carefully taken to the Viventis microscope for time-lapse imaging.

### Single-somite explants

Embryos at the five to seven somite stage obtained from a cross between Utr::mCherry and H2B::GFP heterozygous lines were dechorionated and placed in Leibovitz’s L15 medium (Thermo Fisher, Catalog No. 21083027) in a petridish lined with 2% solidified agarose. A pair of fine forceps (Fine Science Tools, Item No. 11252-00) were used for carefully removing the skin of the embryo. A micro knife (Fine Science Tools, Item No. 10318-14) was then used to scrape off the yolk from the embryo. Two incisions were then made with the knife anterior to the first somite and posterior to the most recently formed somite. This was then followed by an incision along the notochord to separate left and right sides of the embryo. A single somite (somite three or somite four) was then isolated by several careful incisions where the lateral plate mesoderm, neural plate and neighbouring somites were removed. The isolated somite was then transferred to the Viventis imaging chamber using a 20 μl pipette that was pre-coated with Pluronic F-127 (BioVision, Catalog No. 2731) to prevent sticky surfaces. Single somites were imaged with multiple z-slices (2 μm apart) at a frame interval of 2 min for 6 hrs. To determine shape anisotropy of explants, the middle z-plane was chosen and somite contours were first manually segmented in MATLAB. An ellipse was then fit to the segmented 2D contour and the anisotropy in shape, defined as (*b — a*)/(*b + a*), where *b* and *a* are the major and minor axis lengths, was determined. An exponential fit of the change in anisotropy yielded the relaxation time-scale of explants to a spherical shape. For one of the explants, the relaxation to a spherical shape commenced only after an hour after preparation of the explants and therefore for this explant only corresponding data points were fitted.

## Supporting information

Supplemental theory, supplemental figures and supplement movie captions

## Data Availability

Map-projected light sheet data is available for download here: http://doi.org/10.5281/zenodo.4146919

All other data are available upon request from the corresponding author.

## Code Availability

All custom-developed image analysis codes are available for download here: https://github.com/sundar07/SomSeg

## Acknowledgments

We thank members of the Oates lab, M. Gonzales-Gaitan, S. W. Grill, M. Labouesse, P. Tomancak, G. Salbreux, J. Bois, K. Uriu, V. Krishnamurthy, P. Gross, Z. Hadjivasiliou and P. Chugh for comments on the manuscript; fish and bioimaging and optics core facility of École Polytechnique Fédérale de Lausanne. We acknowledge the vital early input of Chen Qian in the project. This work was supported by EPFL, Wellcome (WT098025MA); the Francis Crick Institute, receiving its core funding from Cancer Research UK, the Medical Research Council, and Wellcome; S.R.N. by a Long-Term Human Frontier Science Program postdoctoral fellowship (LT000078/2016); M.P. by the Swiss National Science Foundation Grant No. 200021-165509 and the Simons Foundation Grant No. 454953. S.R.N., M.P., and A.C.O. designed experiments, analyzed data and wrote the paper; S.R.N. performed experiments; M.P. developed theoretical model.

