## Supplemental theory, supplemental figures and supplement movie captions for "Somite surface tension buffers imprecise segment lengths to ensure left-right symmetry"

### Supplementary Information for Somite surface tension buffers imprecise segment lengths to ensure left-right symmetry

#### Mechanical model of AP length adjustment of a newly formed somite

Here we describe the coarse-grained mechanical model of a newly-formed somite. In the model, the somite is considered as a homogeneous cuboid with dimensions $L_{AP}$, $L_{ML}$ and $L_{DV}$ (Fig. 3D(a)) oriented along the main animal axes throughout the relevant time window. In such a cuboid, the velocity gradient tensor is diagonal and the shape changes are related to the corresponding velocity gradient components

$$v_{AP} = \frac{\partial_t L_{AP}}{L_{AP}} \quad (1)$$

$$v_{ML} = \frac{\partial_t L_{ML}}{L_{ML}} \quad (2)$$

$$v_{DV} = \frac{\partial_t L_{DV}}{L_{DV}} \quad (3)$$

Following experimental observation that the volume is conserved, we describe somites as incompressible so that the volume

$$V = L_{AP} L_{ML} L_{DV} \tag{4}$$

is constant. We can express the incompressibility as

$$v_{AP} + v_{ML} + v_{DV} = 0 \tag{5}$$

The shear rate tensor is also diagonal and due to incompressibility, the diagonal components are equal to those of velocity gradients

$$\tilde{v}_{AP} = v_{AP} - \frac{1}{3}\left(v_{AP} + v_{ML} + v_{DV}\right) = v_{AP} \quad , \tag{6}$$

$$\tilde{v}_{ML} = v_{ML} - \frac{1}{3}\left(v_{AP} + v_{ML} + v_{DV}\right) = v_{ML} \quad , \tag{7}$$

$$\tilde{v}_{DV} = v_{DV} - \frac{1}{3}\left(v_{AP} + v_{ML} + v_{DV}\right) = v_{DV} \quad . \tag{8}$$

Stresses acting on somite boundaries arise from somite surface tension $\Gamma$, which originates from a fibronectin-rich extracellular matrix, epithelial cells of the somite and from contact with surrounding tissues. Surface tension stresses acting on each cuboid surface arise from surface tension forces in the neighbouring surfaces acting on the surface boundaries (Sup Fig. 8) and therefore,

$$\sigma_{\Gamma,AP} = -2\Gamma\left(\frac{1}{L_{ML}} + \frac{1}{L_{DV}}\right) \quad , \tag{9}$$

$$\sigma_{\Gamma,ML} = -2\Gamma\left(\frac{1}{L_{AP}} + \frac{1}{L_{DV}}\right) \quad , \tag{10}$$

$$\sigma_{\Gamma,DV} = -2\Gamma\left(\frac{1}{L_{ML}} + \frac{1}{L_{AP}}\right) \quad . \tag{11}$$

PSM ablation experiments established the presence of a compressive stress $\sigma_a$ on the posterior somite surface (Fig. 3C and Sup Fig. 7). Since the PSM is much longer than the somite, any variation of $L_{AP}$ creates only a tiny relative deformation of the PSM. We thus expect that $\sigma_a$

does not depend significantly on $L_{AP}$. Therefore, we consider the PSM to act as a long spring, imposing a dynamic but $L_{AP}$ independent normal stress $\sigma_a$ on the somite posterior surface. To balance the posterior stress $\sigma_a$, we include normal stress $\sigma_a$ on both the anterior and posterior surfaces of the somite (Fig. 3D(c)). To describe the contact between the somite and neural plate on the dorsal side and with yolk on the ventral side, we introduce a time-dependent constraint on $L_{DV} \leq l(t)$, where $l(t)$ is the distance between the yolk and neural plate (Fig. 3D(d)). Convergence-extension flow drives the extension of the somite in the DV direction to fill the available space and the constraint can be written as $L_{DV} = l(t)$, as long as stress between the somite and the neural plate is compressive: $\sigma_{DV} - \sigma_{\Gamma,DV} < 0$, which we discuss below.

As noted in the main text, we do not consider explicitly contact stresses with neighbouring tissues in the mediolateral direction. If such contribution exists it would appear in the model similarly to $\sigma_a$. In the outcome of the model we can consider $\sigma_a$ to account for this contribution as well and not including it explicitly would not change our conclusions.

In the model, we do not consider frictional forces between the somite and its environment and the force balance thus requires stresses in the somite to be constant [1]. We now write the boundary conditions as

$$\sigma_{AP} = \sigma_{\Gamma,AP} + \sigma_a \tag{12}$$

$$\sigma_{ML} = \sigma_{\Gamma,ML} \tag{13}$$

$$L_{DV} = l(t) \tag{14}$$

Developing somites undergo a significant permanent shape change, which suggests that stresses acting on it are beyond the proposed yield stress value [1] and that the somite tissue is in a plastic regime where it flows under stress. Furthermore, our observation of rounding of

[1] The usual compatibility condition is assumed, which ensures that the displacement field remains continuous.

explanted somites (Fig. 3B and Sup Fig. 6) suggests that surface tension stresses are sufficiently above the yield stress value. Taken together, we can write linear constitutive equations relating the somite shear rate and shear stress as

$$\tilde{v}_{AP} = \frac{1}{\eta}\left(\sigma_{AP} + P\right) + \frac{1}{\eta}\zeta_{AP} \quad , \tag{15}$$

$$\tilde{v}_{ML} = \frac{1}{\eta}\left(\sigma_{ML} + P\right) + \frac{1}{\eta}\zeta_{ML} \quad , \tag{16}$$

$$\tilde{v}_{DV} = \frac{1}{\eta}\left(\sigma_{DV} + P\right) + \frac{1}{\eta}\zeta_{DV} \quad , \tag{17}$$

where $\eta$ is somite viscosity and tensor $\boldsymbol{\zeta}$ describes active anisotropic stresses that drive convergence-extension flows [2, 3]. Note that $\boldsymbol{\zeta}$ is traceless: $\zeta_{AP} + \zeta_{ML} + \zeta_{DV} = 0$.

To find the dynamical equation for $L_{AP}$ we subtract Eqs. 15 and 16 and use Eq. 5 to express $v_{ML}$ in terms of $v_{AP}$ and $v_{DV}$. Finally, using the boundary conditions Eqs. 12, 13 and 14, as well as Eq. 1, we obtain

$$\frac{1}{L_{AP}}\partial_t L_{AP} = -\frac{1}{2l(t)}\partial_t l(t) + \frac{1}{2\eta}\sigma_a(t) + \frac{1}{2\eta}\left(\zeta_{AP} - \zeta_{ML}\right) - \frac{\Gamma}{\eta}\left(\frac{l(t)L_{AP}}{V} - \frac{1}{L_{AP}}\right) \tag{18}$$

Solving this equation would require knowledge of time dependence of $l(t)$, contact stress $\sigma_a(t)$ and possibly active stress tensor $\zeta$. However, in this work we are interested in somite size adjustment and an extensive study of how $L^0_{AP}$ is set is beyond our current scope. Therefore, we assume that a well defined value of $L^0_{AP}$ exists, as observed in experiments, which we approximate as a constant in time defined by a combination of time-varying contributions

$$L^0_{AP} = \frac{V\eta}{2\Gamma l(t)}\left(\sqrt{s(t)^2 + \frac{4\Gamma^2}{\eta^2}\frac{l(t)}{V}} + s(t)\right) \quad , \tag{19}$$

where $s(t) = -\partial_t l(t)/(2l(t)) + \sigma_a(t)/(2\eta) + (\zeta_{AP} - \zeta_{ML})/2$. Now, given a constant value of $L^0_{AP}$, we proceed to test its stability by considering a small perturbation $\delta L_{AP} = L_{AP} - L^0_{AP}$.

To the lowest order, the perturbation follows

$$\partial_t \delta L_{AP} = -\frac{\Gamma}{L^0_{AP}\eta}\left(1 + \frac{(L^0_{AP})^2 l(t)}{V}\right)\delta L_{AP} \quad . \tag{20}$$

Therefore, any variation of $L_{AP}$ from $L^0_{AP}$ will be reduced in time. The relaxation rate is time-dependent through $l(t)$. However, for observed typical values of $L_{DV}(0) \approx 39\ \mu m$, $L_{DV}(1hr) \approx 53\ \mu m$, $V \approx 1.2 \cdot 10^5\ \mu m^3$, and $L^0_{AP} \approx 53\ \mu m$, the variable term changes from $0.91$ at initial time to $1.24$ at $1\ hr$. Therefore, the relaxation rate changes by about $16\%$ and for simplicity we approximate it with its mean value

$$\partial_t \delta L_{AP} \approx -\frac{2.1\Gamma}{L^0_{AP}\eta}\delta L_{AP} \quad . \tag{21}$$

The relaxation time-scale is then

$$\tau \approx \frac{\eta L^0_{AP}}{2.1\Gamma} \tag{22}$$

In experiments the somite AP length is measured at the time of formation and at 1 and 2 hours later. In order to determine the time-scale $\tau$ from experimental data we write the Eq. 21 as

$$\partial_t L_{AP} = -\frac{1}{\tau}\left(L_{AP} - L^0_{AP}\right) \quad , \tag{23}$$

which can be solved for

$$L_{AP}(t) = e^{-\frac{t}{\tau}} L_{AP}(0) + \left(1 - e^{-\frac{t}{\tau}}\right) L^0_{AP} \quad . \tag{24}$$

Therefore, the time-scale $\tau$ can be determined by performing a linear fit to the experimental data $L_{AP}(1hr)$ vs. $L_{AP}(0hr)$ and $L_{AP}(2hr)$ vs. $L_{AP}(0hr)$. Then, $\tau = -1/\ln a$ where $a$ is the slope coefficient of the linear fit. It is interesting to point out that even in a more general case where $L^0_{AP}$ is not constant but relaxes from initial value at the moment of the somite formation to a final equilibrium value this procedure still provides a good measure of $\tau$. Namely, for $L^0_{AP}(t) = L_0 + \Delta e^{-t/\tau_0}$ dynamics of somite $AP$ length follows

$$L_{AP}(t) = e^{-\frac{t}{\tau}} L_{AP}(0) + \left(1 - e^{-\frac{t}{\tau}}\right) L_0 + \left(e^{-\frac{t}{\tau_0}} - e^{-\frac{t}{\tau}}\right) \Delta \quad , \tag{25}$$

which gives the same coefficient of proportionality between $L_{AP}(t)$ and $L_{AP}(0)$.

Explanted somites relax over time from an irregular shape to a sphere. We quantified the relaxation time-scale of the explants $\tau_e$, as shown in the main text. In order to compare $\tau_e$ with relaxation time-scale $\tau$ of an *in vivo* somite, we approximate the former with an analytical expression for relaxation of a fluid prolate ellipsoid under surface tension [4]

$$\tau_e \approx \frac{19}{20} \frac{\eta R}{\Gamma} \quad , \tag{26}$$

where $R$ is the final radius of the sphere and viscosity of the surrounding solution is neglected in comparison to the tissue viscosity $\eta$. For the mean volume estimate mentioned above, we find $R = \sqrt[3]{3V/(4\pi)} \approx 31\ \mu m$ so that $\tau_e \approx 29\eta/\Gamma\ \mu m$. Using the measured value $\tau_e = 1.1 \pm 0.4\ hr$ we can estimate $\eta/\Gamma \approx 0.04 \pm 0.01\ hr/\mu m$. For *in vivo* somites, we use Eq. 22 to quantify the same quantity $\eta/\Gamma \approx 0.063 \pm 0.004\ hr/\mu m$. Although not exactly the same the two measured values are similar, which suggests that surface tension is indeed a major contributor to somite size robustness. Differences between *in vivo* and explant values could stem from additional viscous and/or frictional dissipation occurring *in vivo*.

Finally, we note that the condition $\sigma_{DV} - \sigma_{\Gamma,DV} < 0$, required for the DV constraint $L_{DV} = l(t)$ to be fulfilled, can be expressed as

$$\frac{3}{2}\eta \left(\frac{1}{L_{DV}} \partial_t L_{DV}\right) - \frac{3}{2}\zeta_{DV} - 2\Gamma \frac{1}{L_{DV}} + \Gamma \frac{1}{L_{AP}} + \Gamma \frac{1}{L_{ML}} < 0 \tag{27}$$

The somites that we observe remain in contact with the neural plate during the experiment. Over that time $L_{DV}$ grows and typically overcomes both $L_{AP}$ and $L_{ML}$, which suggest that $q_{DV}$ is positive and sufficiently large to maintain this condition. It is interesting to further remark that in the absence of mechanical constraints on the DV surfaces, we would have $\sigma_{DV} = \sigma_{\Gamma,DV}$ and the dynamics of DV somite dimension would be determined by equating left hand side of Eq. 27 with $0$, which could be relevant for description of tail somites that may have no mechanical contacts at DV boundaries.

### Supplementary Movie Captions

**Supplementary Movie 1: Multiview movies** Zebrafish embryos were imaged from 6 angles (as indicated) using Zeiss Z1 and the maximum intensity projection of nuclei channel (H2B::GFP) in each angle is shown.

**Supplementary Movie 2: Map projection allows visualization and analysis of left-right somite morphogenesis simultaneously** Multiview acquisitions were fused and map projected. Top, map projected time-lapse of nuclei channel (H2B::GFP); bottom, corresponding utrophin channel (Utr::mCherry). Anterior is to the left and posterior is to the right. Single map projected layers from each time point were chosen and shown here.

**Supplementary Movie 3: Single-somite explants round up over time** Single somites were isolated and cultured *in vitro*. A single z-slice from each time point was chosen and shown here.

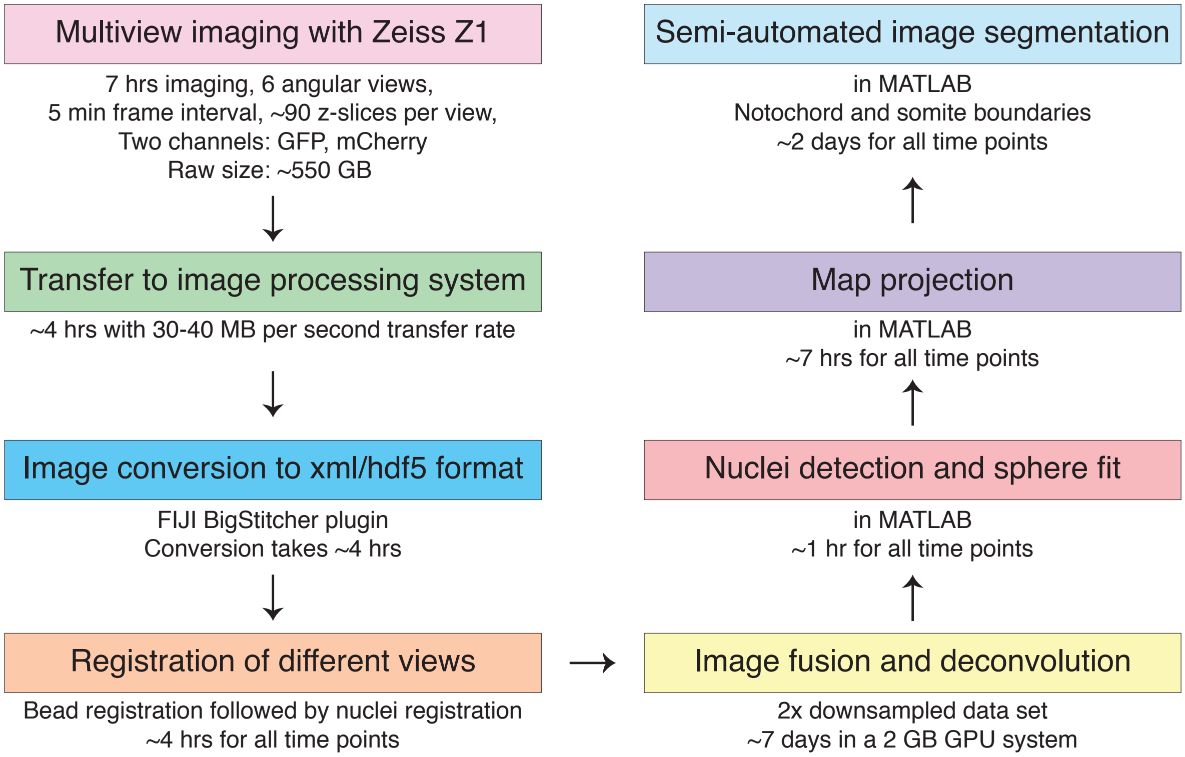


Sup Figure 1: **Steps in multiview data acquisition and processing** The boxes represent the major steps involved and below each box, respective details are included.

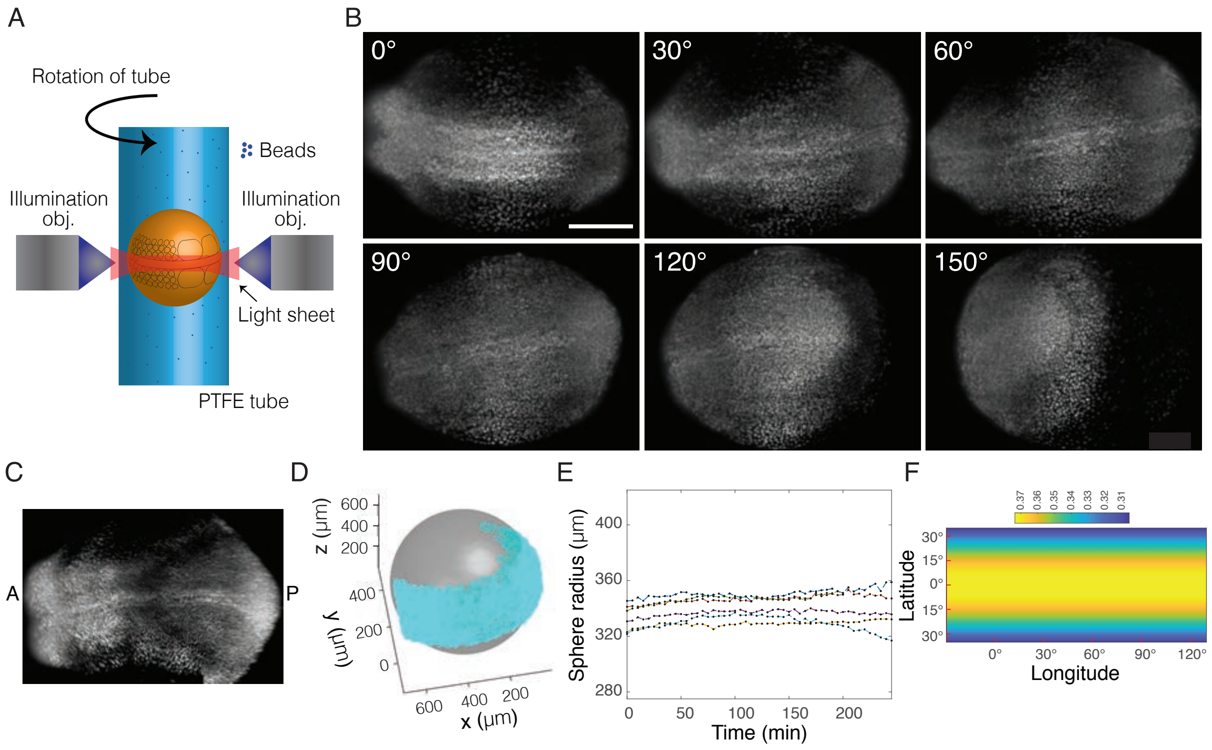


Sup Figure 2: **Multiview data acquisition and processing** (A) Schematic of multiview imaging setup. A zebrafish embryo in its chorion is mounted in a PTFE tube filled with 0.25% agarose and fluorescent beads. An embryo with its AP axis oriented along the circumference of the tube, as indicated, is chosen for multiview imaging. Rotation of the tube allows imaging the embryo from different angles. (B) The first time point of a histone transgenic line viewed from 6 angles. Images represent respective maximum intensity projections. Scale bar, 100 μm. (C) Representative image of multiview fusion, which was performed using the MultiView Reconstruction FIJI plugin. (D) Sphere fit (in gray) of a representative fused image. Cyan dots represent individual nuclei. (E) The radius of the sphere obtained from the fit does not change over the analysis time window. Individual lines represent the 6 multiview movies used for analysis. (F) Heat map of change in pixel size in equidistant cylindrical map projection. Note that the change in pixel size is negligible within 15° from the equator, which corresponds to regions of somite formation (compare with Fig. 1A).

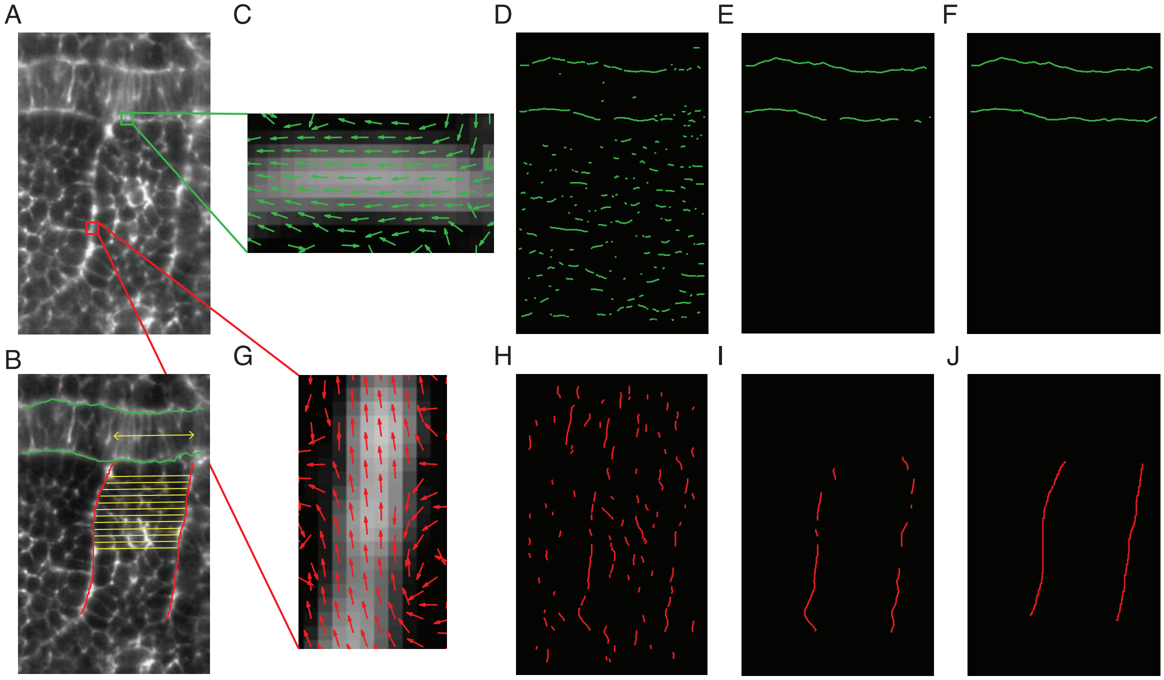


Sup Figure 3: **Steps in custom-developed boundary segmentation method** (A) Raw zoomed in image of a representative region of an embryo imaged using Viventis microscope. (B) Image in (A) overlaid with segmented notochord (in green) and somite boundaries (in red). Segment length (indicated in yellow) is determined with respect to the notochord. Yellow arrow represents local angle of the notochord. (C), (G) Quiver plots of the eigen vectors upon application of Frangi vesselness filter. Orientation of the vectors represent directions of least change in fluorescent intensity. Along the notochord, these directions are horizontal (C), while along somite boundaries, they are vertical (G). (D), (H) Raw image in (A) is subdivided into two parts based on the directions of the eigen vectors, thus leading to images with horizontal (D) or vertical lines (H). (E), (I) Lines that are less than 15 pixels apart are joined together by straight lines following which lines that do not represent notochord or somite boundaries are discarded. (F), (J) Any remaining gaps in boundary detection are then connected by straight lines.

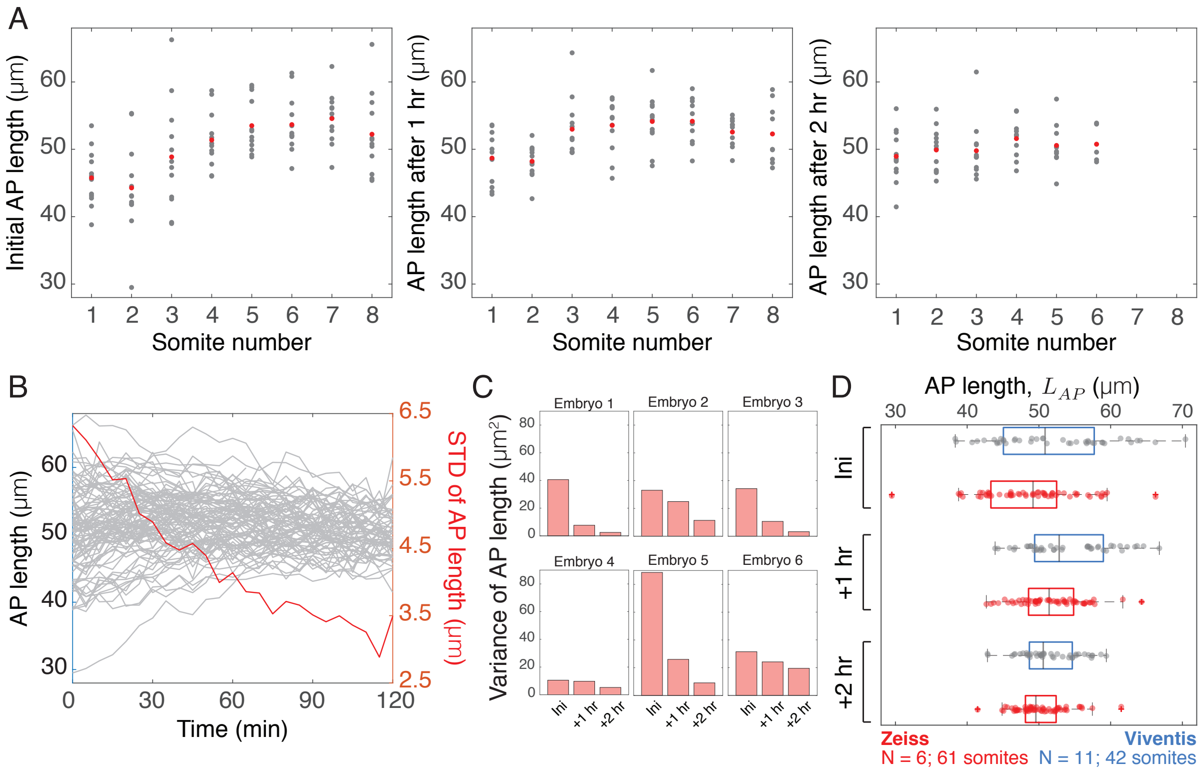


Sup Figure 4: **AP lengths of anterior somites reduce variability over time** (A) Initial AP lengths, lengths at 1 hr and lengths at 2 hr of somites one to eight are shown across the six multiview data sets. Both left and right somites are included in the same plot. Gray dots represent individual somites. Red dots, mean AP length. Somites seven and eight were not segmented at two-hour time points. (B) Change in AP length over time for all eight segmented left-right somites across six embryos. Note the decrease in standard deviation (red) over time. (C) Variance of AP lengths over time in each of the six embryos imaged with Zeiss Z1. (D) Comparison of AP lengths over time across zeiss and viventis data sets.

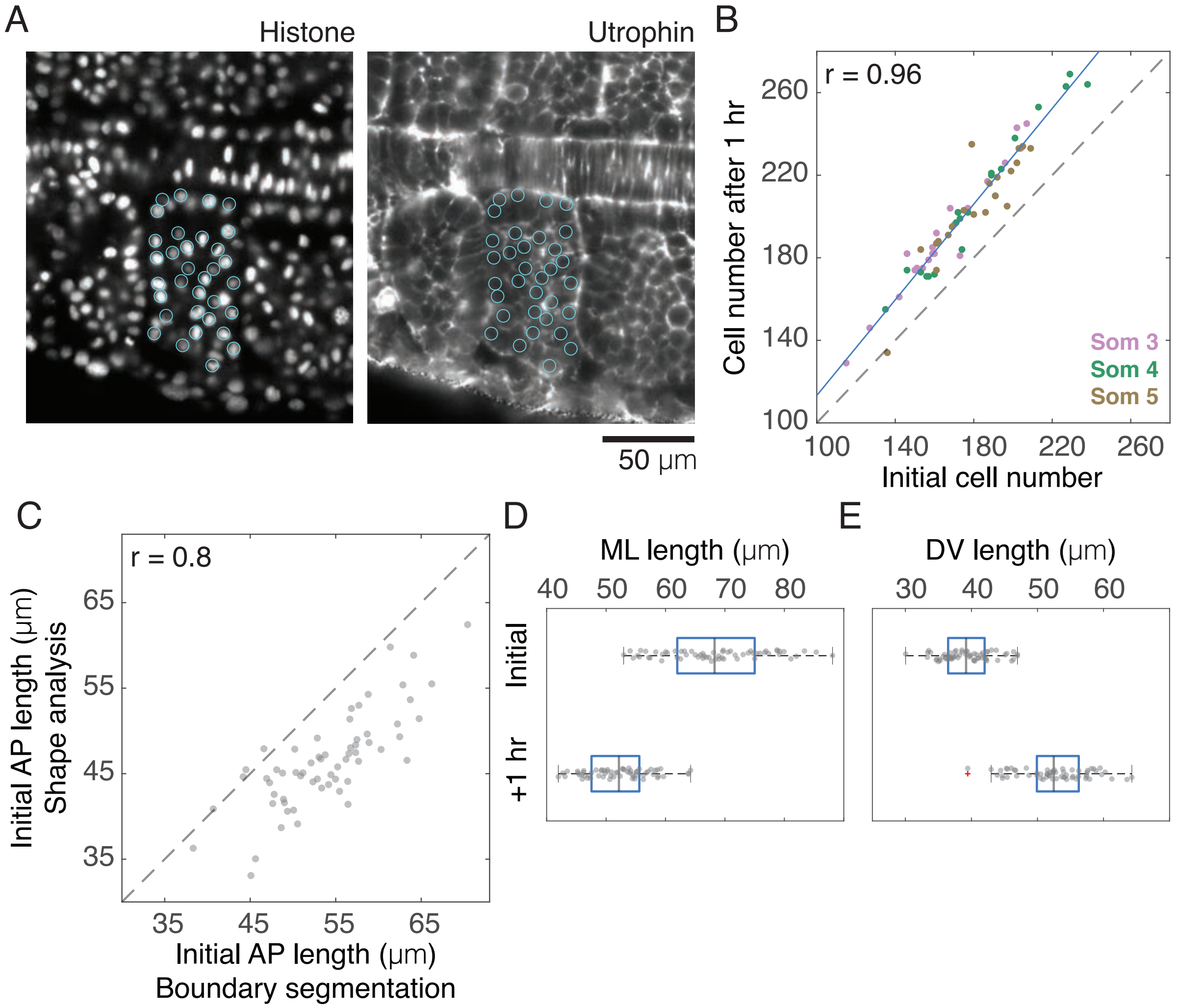


Sup Figure 5: **Somite shape analysis** (A) Snapshots from Mastodon, a FIJI plugin, indicating detected nuclei (cyan circles) in corresponding histone and utrophin fluorescent images. A single zoomed in z-slice of an embryo imaged using Viventis microscope is shown. (B) Comparison of initial and one-hour cell numbers of somites (r, 0.96 [0.93,0.98]). Somites three to five are separately colored. Dashed line, slope=1 in (B,C); blue, regression line. (C) Positive correlation of somite AP lengths obtained from boundary segmentation method and shape analysis (r, 0.8 [0.69,0.87]). Because nuclei are used for determining lengths in the shape analysis, AP lengths are consistently smaller in the shape analysis compared to lengths determined from boundary segmentation. (D-E) Comparison of initial and one-hour ML (D) and DV lengths (E) indicate decrease and increase in lengths respectively.

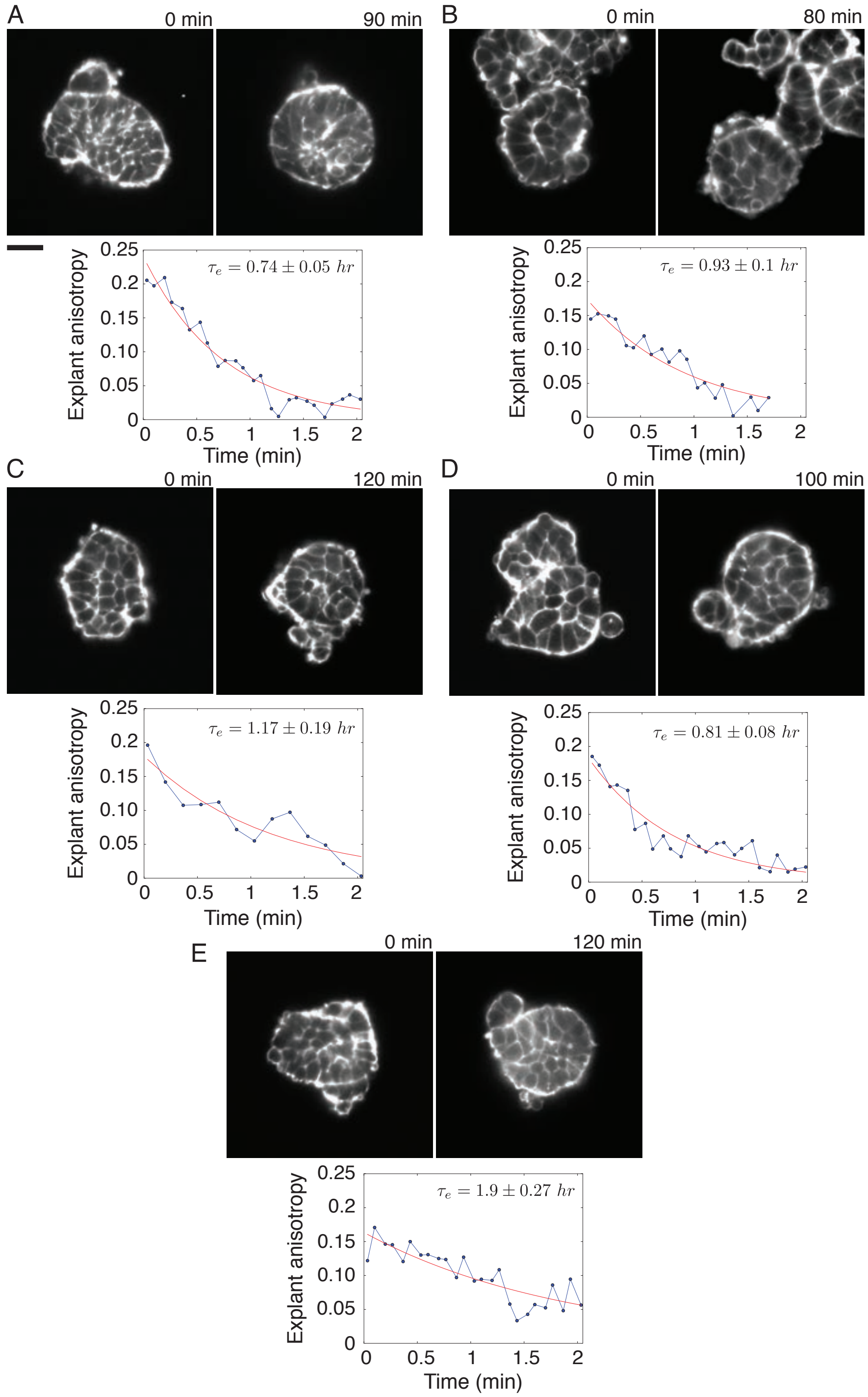


Sup Figure 6: **Shape change in single-somite explants** Left image in each of the five examples (A) to (E) shows initial shape of explants. Right images represent time points when explants acquired spherical shape. Scale bar, 25 μm. (A) is same as shown in Fig. 3B. An exponential fit of explant anisotropy over time is shown below each example. Respective relaxation time scales, $\tau_e$ are as indicated.

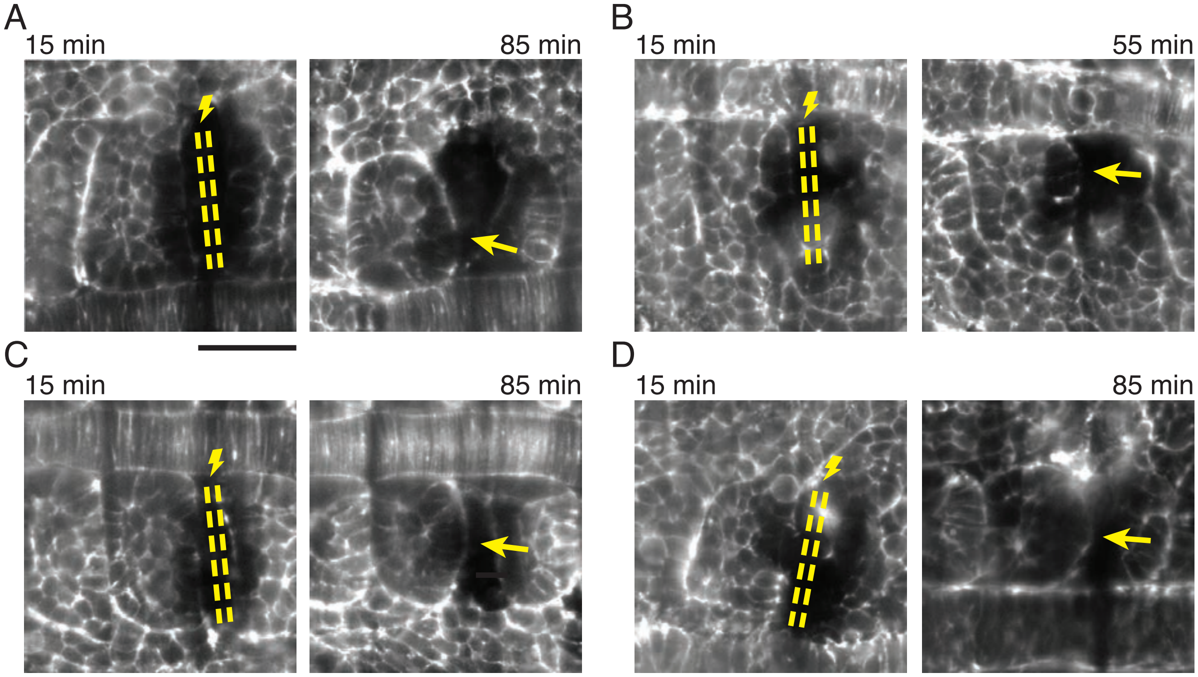


Sup Figure 7: **Ablation of presomitic mesoderm** Representative ablation examples (A) to (D) in addition to the example shown in Fig. 3C. In the left image in each example, yellow lines indicate ablation of presomitic mesoderm adjacent to most recently formed boundary. Imaging in Viventis microscope was started 15 min after ablation. Right image in each example shows a prominent bulge of the boundary (arrow). Scale bar, 50 μm.

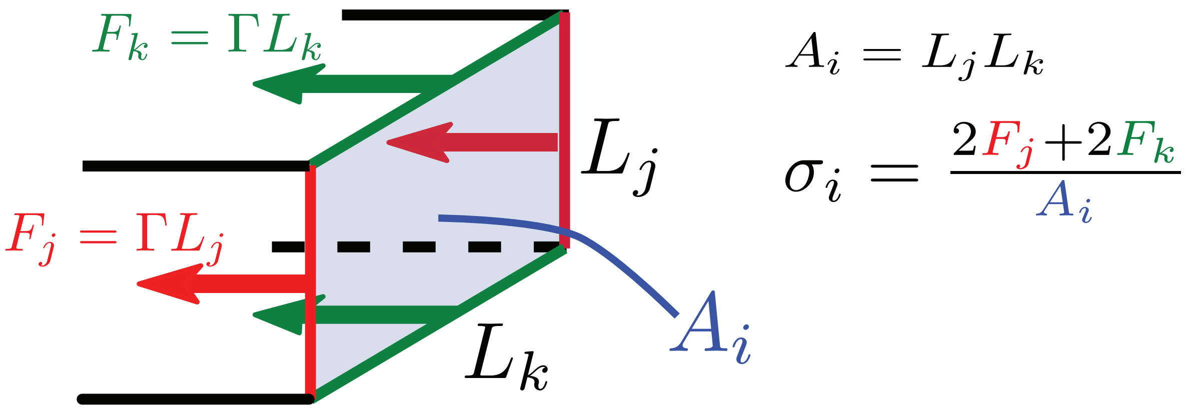


Sup Figure 8: **Normal stress due to surface tension on surface.** Surface tension on the surfaces of the model somite exerts normal stress on the neighbouring surfaces. Surface tension forces $F_j$ (red) and $F_k$ (green) acting on the light blue surface $i$ are proportional to the surface edge lengths $L_j$ and $L_k$. The normal stress $\sigma_i$ is then normal force per surface area $A_i$ from which we obtain Eqs. 9, 10 and 11.

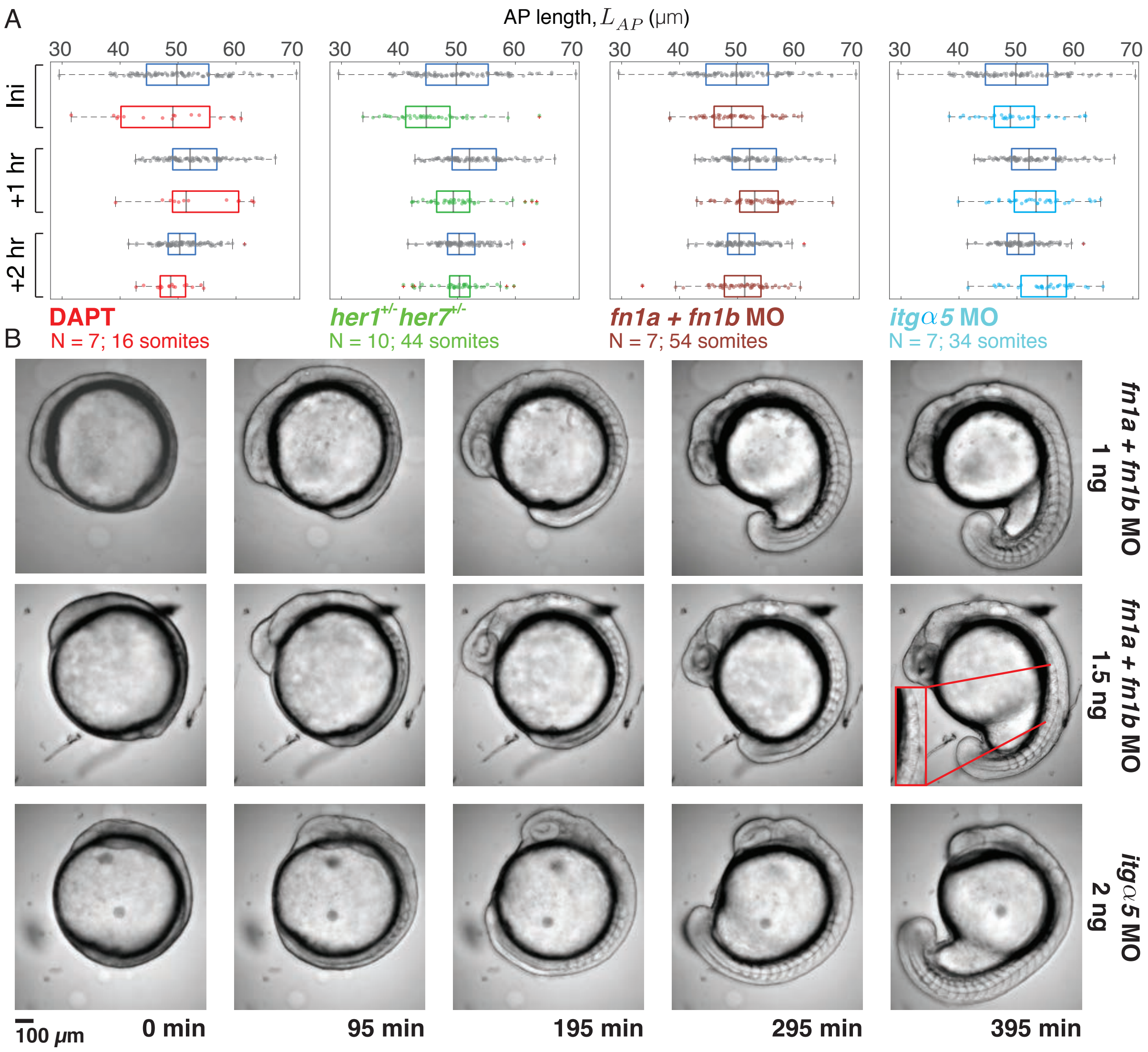


Sup Figure 9: **Integrin and Fibronectin, molecules involved in surface tension, facilitate length adjustment.** (A) Comparison of AP length $L_{AP}$ in *her1;her7* heterozygous mutants, DAPT treated embryos, *integrinα5* and *fibronectin1a* and *1b* MO injected embryos. (B) Injection of 1.5 ng of *fn1a* and *fn1b* MOs results in disintegration of anterior somites by the $10^{th}$ somite stage (zoomed in inset in red at 395 min), while anterior somites remain intact when 1 ng injection is performed. Snapshots of *itgα5* MO (2 ng) injected embryo.

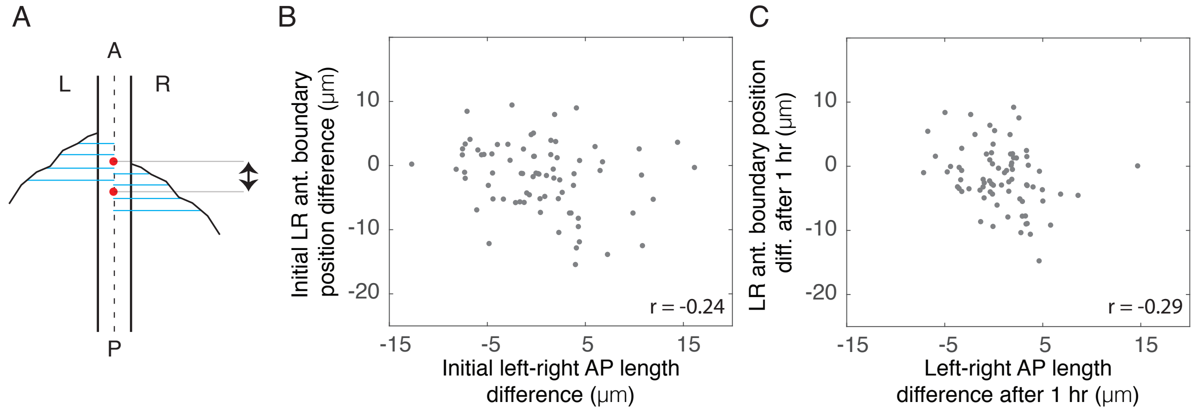


Sup Figure 10: **Left-right difference in somite length is weakly correlated with anterior somite boundary difference.** (A) Schematic of measurement of boundary position difference. The points of intersection of orthogonal lines (cyan) from somite boundaries on the left (L) and right (R) sides with the middle of the notochord (black dashed line) are determined. A difference in the median position (red dots) of these points of intersection represents boundary position difference (black arrow). Boundary positions increase positively from P to A. (B-C) AP length difference between left-right somite pairs is weakly correlated with anterior boundary position difference both initially (r, -0.24 [-0.45,-0.03]) (B) and after 1 hour (r, -0.29 [-0.48,-0.1]) (C).
